# HIV reservoir size is determined prior to ART initiation and linked to CD8 T cell activation and memory expansion

**DOI:** 10.1101/580308

**Authors:** Genevieve E Martin, Matthew Pace, Freya Shearer, Eva Zilber, Jacob Hurst, Jodi Meyerowitz, John P Thornhill, Julianne Lwanga, Helen Brown, Nicola Robinson, Emily Hopkins, Natalia Olejniczak, Nneka Nwokolo, Julie Fox, Sarah Fidler, Christian B Willberg, John Frater, on behalf of the CHERUB investigators

## Abstract

Initiation of antiretroviral therapy (ART) in early compared with chronic HIV infection is associated with a smaller HIV reservoir. This longitudinal analysis of 63 individuals who commenced ART during primary HIV infection (PHI) investigates which pre-and post-therapy factors associate most closely with reservoir size (HIV DNA) following treatment initiation during PHI. The best predictor of reservoir size at one-year was pre-ART HIV DNA which was in turn significantly associated with CD8 memory differentiation (effector memory, naïve and T-bet^neg^Eomes^neg^ subsets), CD8 T cell activation (CD38 expression) and PD-1 and Tim-3 expression on memory CD4 T cells. No associations were found for any immunological variables following one-year of ART. HIV reservoir size is determined around the time of ART initiation in individuals treated during PHI. CD8 T cell activation and memory expansion are linked to HIV reservoir size, suggesting the importance of the initial host-viral interplay in eventual reservoir size.

## Introduction

HIV persists despite long-term suppressive antiretroviral therapy (ART) in a reservoir of latently infected cells (1-3). Reducing the size of this reservoir is a focus of curative interventions as a smaller HIV reservoir is associated with delayed clinical progression and viral rebound following treatment interruption (4-6).

While there are only a few studies that explore the longitudinal formation of the HIV reservoir, there is some evidence that T cell immunity prior to the start of ART may be key. In the SPARTAC trial, both pre-therapy CD4 and CD8 T cell responses, as well as protective HLA class I alleles, were linked with lower levels of HIV DNA (7). Additionally, recent work from a different cohort treated during acute infection showed that HIV-specific T cell responses could reduce viral load (VL), and limit reservoir seeding, when antiretroviral therapy (ART) was initiated at peak viraemia (8).

T cell activation – a mediator of disease progression and non-AIDS co-morbidities (9-12) – has also been shown in cross-sectional studies to be related to HIV reservoir size (7, 13-15). Two recent longitudinal studies aimed to clarify this relationship. In individuals who initiated ART during chronic infection, no link between pre-ART inflammation/activation was observed with reservoir size over time (16). Contrasting with this, the finding that several soluble biomarkers in acute infection, including many linked to interferon-α (IFN-α) signalling, were identified as correlates of subsequent reservoir size (17) suggests a link between early immune activation and the formation of the reservoir.

Immune checkpoint receptor (ICR) expression has also been linked to the size of the HIV reservoir (7, 13, 14, 18, 19). A limitation of many of these studies is that ICR expression has often been measured on bulk T cells. The HIV reservoir is preferentially found in memory subsets (19-22) which also express higher levels of ICRs (18, 23-26). As such, ICR expression on bulk T cell subsets could be a surrogate for altered memory distribution and a potential confounder (27). Furthermore, many of these relationships were demonstrated for a single marker, even though the expression of many ICRs is closely related to one another – and also to T cell activation (23-26, 28-31).

The ‘HIV Reservoir Targeting with Early Antiretroviral Therapy’ (HEATHER) cohort is a prospective study of individuals commencing ART during primary HIV infection (PHI), and provides the opportunity to study reservoir dynamics over time in well-characterised participants. Here, we define the relationship of several key immunological and clinical parameters with the size of the HIV reservoir and show that the key events in setting the reservoir occur early, prior to starting ART.

## Results

### Baseline clinical characteristics

We studied 63 individuals enrolled in HEATHER, a longitudinal observational cohort of more than 300 participants who commenced ART during PHI. Individuals were selected based on availability of samples and access to clinical records at the time of the study. Clinical and demographic details are listed in Table 1. All were male and had a median age of 36 (IQR 28-41) years at the time of ART start. They commenced ART a median of 29 (IQR 14-46) days following a confirmed HIV diagnosis and a median of 49 (IQR 33-90) days following estimated seroconversion. Different methods for diagnosing PHI (Table 1) were used; 26 participants (41%) were P24 antigen positive without detectable antibodies, consistent with Fiebig stages I or II at the time of diagnosis.

**Table 1.** Demographic and clinical characteristics of participants

|  |  |
| --- | --- |
| Sex |  |
| • Male | 63 (100%) |
| Age (years) | 36 [28 - 41] |
| Time between date of confirmed HIV+ test and ART initiation (days) | 29 [14 - 46] |
| Time between estimated date of seroconversion and ART initiation (days) | 49 [33 - 90] |
| Time between ART initiation and first VL <50 copies/ml (days)<sup>A</sup> | 133 [93 - 241] |
| Baseline CD4 T cell count (cells/μL)<sup>B</sup> | 527 [406 - 645] |
| Baseline CD8 T cell count (cells/μL)<sup>B</sup> | 1033 [831 - 1326] |
| Baseline CD4 to CD8 ratio<sup>B</sup> | 0.52 [0.37 – 0.76] |
| Baseline viral load (log<sub>10</sub> copies[HIV RNA]/ml) | 5.5 [4.6 – 6.5] |
| Method for diagnosing primary HIV infection |  |
| • Antigen positive (p24 or PCR) but antibody negative | 26 (41%) |
| • Rising antibody titre | 1 (1.5%) |
| • Negative test within 6 months of positive test | 30 (48%) |
| • Recent incidence testing algorithm | 6 (9.5%) |
| Mode of acquisition |  |
| • MSM | 56 (89%) |
| • MSW | 1 (1.5%) |
| • Unknown/unrecorded | 6 (9.5%) |
| Initial ART regimen |  |
| • Unknown/unrecorded | 3 (4.8%) |
| *Backbone* |  |
| • tenofovir containing | 55 (87%) |
| • abacavir containing | 5 (7.9%) |
| *Additional agent(s)* |  |
| • protease inhibitor | 36 (57%) |
| • NNRTI | 11 (17%) |
| • integrase inhibitor | 12 (19%) |
| • protease inhibitor + integrase inhibitor | 1 (1.5%) |

Demographic and baseline clinical characteristics of included participants from the HEATHER cohort. Values given represent n (%) for categorical variables and median (interquartile range) for continuous variables. Abbreviations: MSM, men who have sex with men; MSW, men who have sex with women, NNRTI, non-nucleoside reverse-transcriptase inhibitor. <sup>A</sup> 60/63 individuals were virologically suppressed to <50 copies/ml by 1 year visit <sup>B</sup> Data available for 62/63 individuals

The participants had a high median baseline VL (5.5 log_10_ copies/ml; IQR 4.6-6.5), which declined on ART (Fig. 1A). 60/63 (95%) were virologically suppressed (<50 copies/ml) after one year of ART (the frequency of viral load sampling and time to suppression is shown in Supplementary Fig. 1). The three individuals who had not achieved VL<50 copies/ml by one year had estimated VLs (average of those either side of the 1 year time-point) of 40, 76 and 190 copies/ml, and achieved undetectability shortly after. The dynamics of CD4 and CD8 count, as well as CD4 to CD8 ratio following ART initiation are shown in Fig. 1B-D.

**Figure 1.**
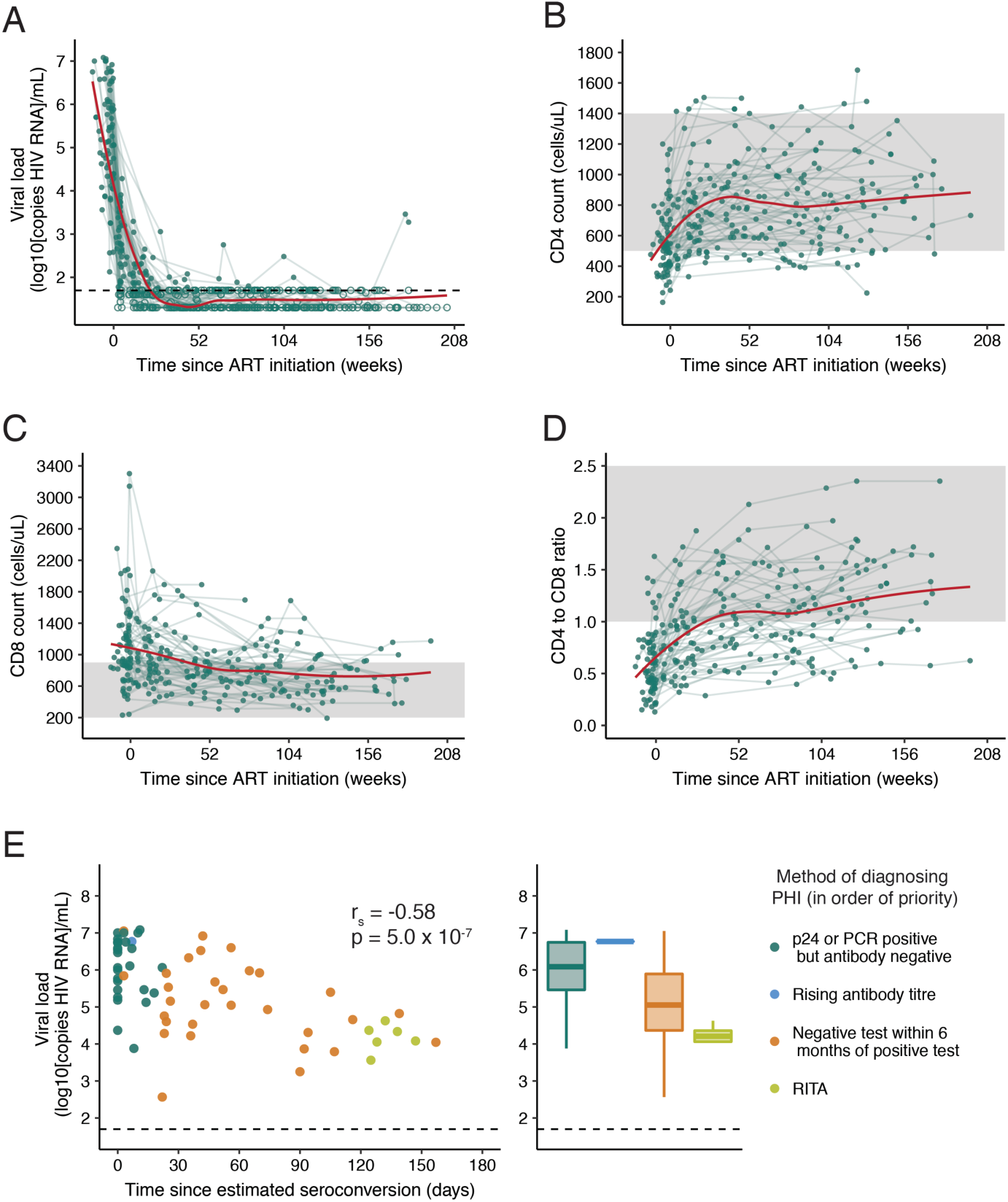
Measures of clinical progression during treated primary HIV infection. **(A)** Viral load (VL), **(B)** CD4 count **(C)** CD8 count and **(D)** CD4 to CD8 ratio in the 4 years following ART initiation (n=63). A trend line (shown in red) has been fitted using local polynomial regression fitting (LOESS) smoothing with an α value of 0.75. For **(A)** exact values are shown as closed circles, and those below the limit of detection are shown as open circles; the black dashed line indicates 50 copies/ml. In **(B-D)** the shaded region shows the normal range for these parameters. **(E)** Baseline VL relative to the number of days this was measured after estimated seroconversion (left panel; Spearman’s correlation) with the same data as histograms stratified by method of diagnosing PHI (right panel). If individuals met multiple diagnostic criteria, they are plotted as the criterion with the most reliability in estimating date of seroconversion; these are shown in the legend in the order of priority used.

There was a relationship between the first measured (or ‘baseline’) pVL and the method used to diagnose PHI (Fig. 1E). Baseline VL was higher when measured closer to estimated seroconversion (r_s_=-0.58, p=5.0×10^-7^), suggesting that viral load is of limited utility as a predictive variable in PHI compared with chronic HIV infection, as a stable ‘set point’ has not yet been reached.

### HIV reservoir size after one year of ART is predicted by pre-ART HIV DNA levels

Quantitation of total HIV DNA (copies per 10^6^ CD4 T cells) is used here as the surrogate measure of reservoir size on ART. Other measures (e.g. integrated HIV DNA, cell-associated RNA, quantitative viral outgrowth) were not used for this analysis due to the high numbers of predictive variables being tested. Of note, the term “reservoir” is only used for HIV DNA measurements on suppressive ART as pre-ART DNA measurements will capture cells with active viral replication. Compared with pre-ART levels, HIV DNA decreased a mean of 0.9 log_10_ copies following one year of therapy (Fig. 2A; p=2.2×10^-16^). HIV DNA levels pre-therapy and following one year of ART were highly correlated (Fig. 2B; r=0.76, p=1.2×10^-12^). For a subset of 18 individuals, levels of total HIV DNA were also available three years post-ART initiation, and had declined a further 0.3 log_10_ copies since year one. (HIV DNA levels were also correlated between those two measurements (Supplementary Fig. 2; r-0.57, p=0.013)).

**Figure 2.**
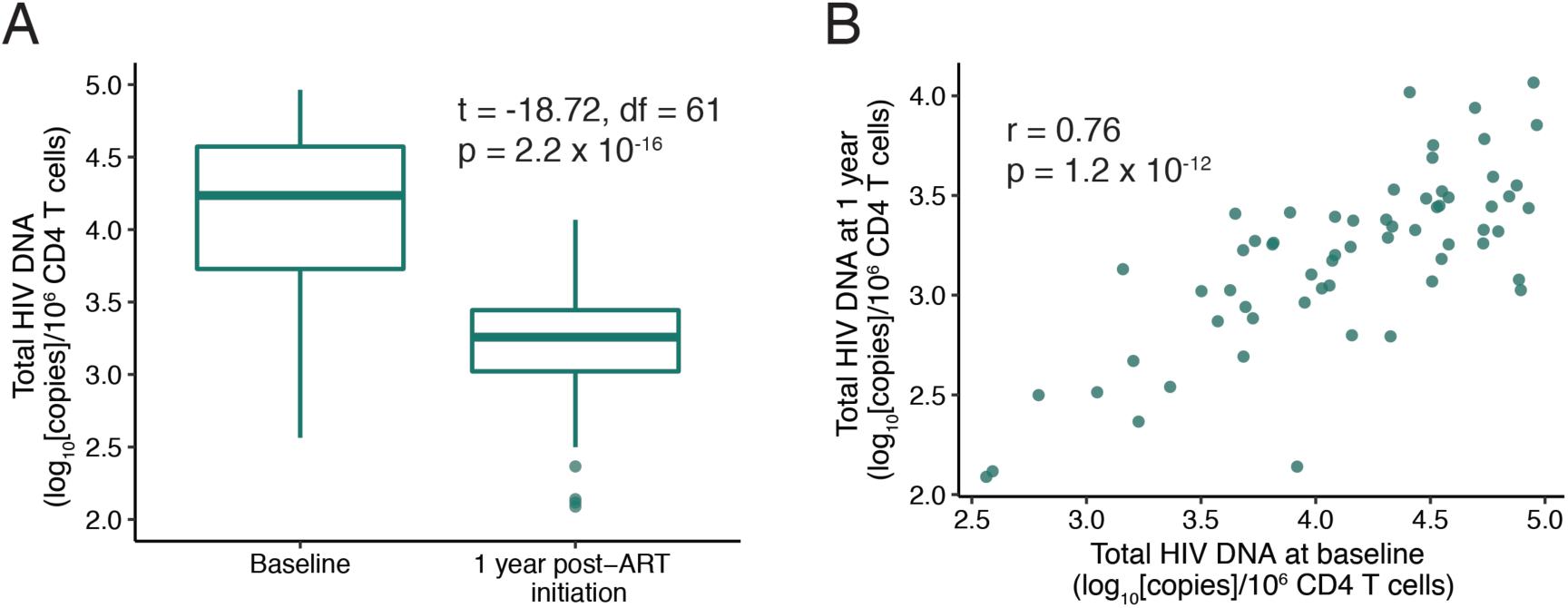
Total HIV DNA during treated primary HIV infection. **(A-B)** Relationship between total HIV DNA measured at baseline and 1 year following ART initiation (n=62). For **(A)** comparison was made using a paired t-test and in **(B)** a Pearson’s correlation was performed.

### Immunological and clinical variables associated with HIV DNA are highly correlated

We next determined which clinical and immunological variables were most closely associated with the size of the HIV reservoir. Clinical variables included CD4 count, CD8 count, VL, CD4:CD8 ratio, time to ART start, and time to VL suppression on ART. Immunological measures included flow cytometric quantitation (Fig. 3A) of CD4 and CD8 memory subsets, CD38 expression, ICR expression (PD-1, TIGIT, Tim-3 on memory CD4 and CD8 T cells), the proportion of T-bet/Eomes expressing CD8 populations (32) and soluble plasma ICRs (sPD-1 and sTim-3).

**Figure 3.**
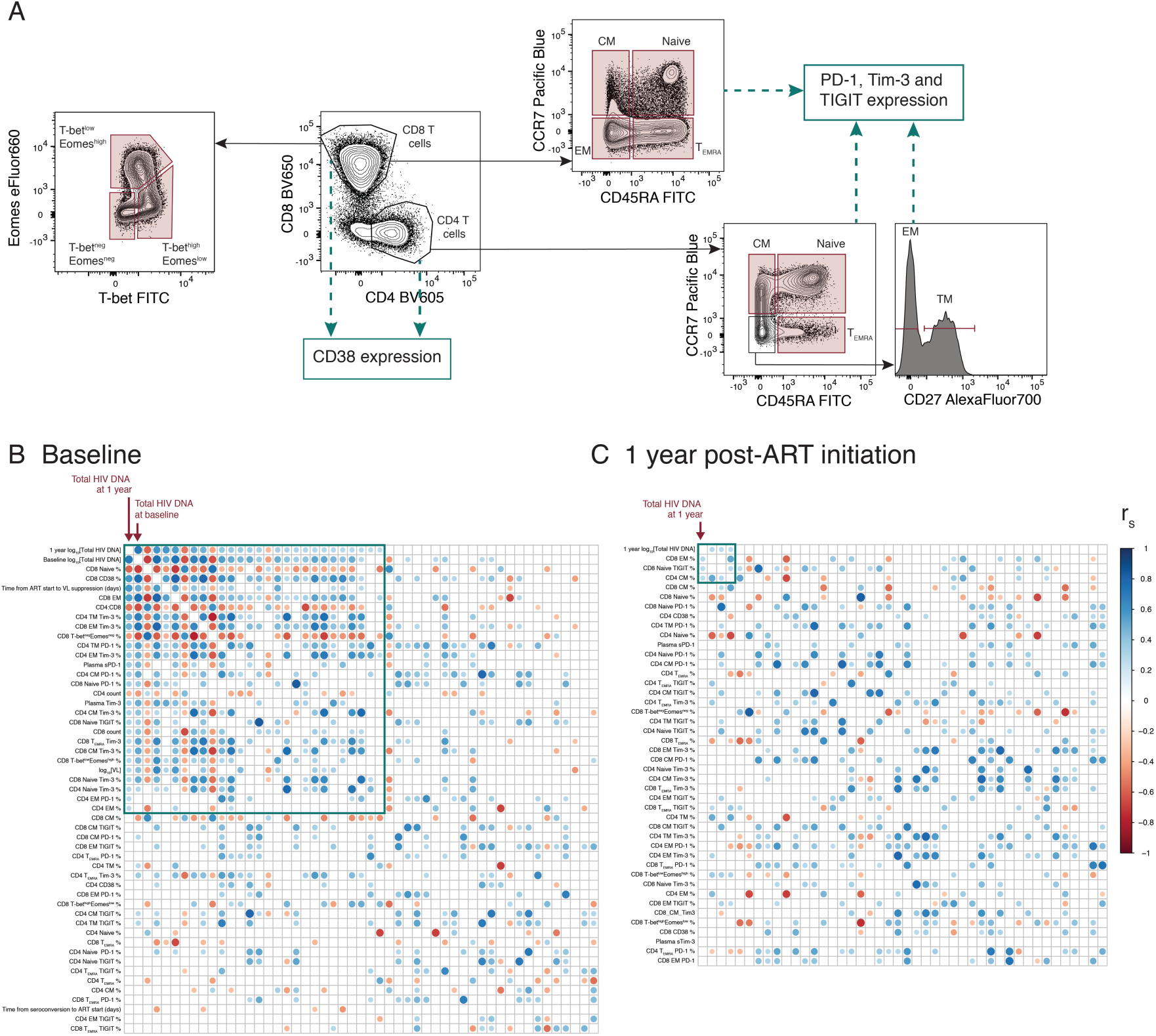
Immunologic and clinical variables associated with HIV reservoir size are highly correlated with one another. **(A)** Schematic showing the T cell subsets and surface markers measured by flow cytometry in this analysis. The frequency of populations gated in red was included in analysis, as well as the expression of CD38, PD-1, Tim-3 and TIGIT on populations marked. **(B-C)** shows the correlations between clinical/immunological variables and HIV reservoir size. Corrplots showing the relationship between HIV reservoir size at 1 year (log_10_ [Total HIV DNA]) and immunological/clinical variables (n=63) measured at baseline **(B)** or following 1 year of ART **(C)**. The same immunological variables were included at both time points, and clinical variables at baseline only. Reservoir size at 1 year (1 year log_10_[Total HIV DNA]) is shown in the top left corner and is marked. For both B and C, variables have been ranked based on the magnitude of absolute correlation coefficient with 1 year log_10_[Total HIV DNA] in decreasing order from the top left hand corner. The size and colour of each circle corresponds to the correlation coefficient between any two variables. Correlation coefficients were calculated using the Spearman method with pairwise complete observations; only correlations significant at a 0.05 level are shown (other boxes are left blank). Green boxes enclose variables which have a significant correlation with 1 year log_10_ [Total HIV DNA] at a 0.05 level. Abbreviations: CM, central memory; EM, effector memory.

Several parameters were highly correlated with HIV DNA. Corrgrams were used as a screen to explore the relationship of variables measured prior to ART initiation (baseline; Fig. 3B) and after one year of ART (Fig. 3C), with the HIV reservoir at one year. Each row or column in the corrgram represents a different variable ordered by the strength of the correlation with reservoir size at one year (in the top left corner). A circle indicates a correlation between two variables (p<0.05), the size and intensity corresponding to the magnitude of the correlation coefficient. Variables with a statistically significant relationship with reservoir size at 1 year are marked by a green square.

Two important observations can be made. Firstly, the corrgrams for variables at baseline and one year look very different (Figs. 3B,C). When exploring variables measured immediately prior to ART initiation, we can see that those variables which were most closely related to reservoir size (top left hand corner) were also highly correlated with each other. Not surprisingly, the variable with the strongest correlation with HIV reservoir size at one year was the level of HIV DNA measured at baseline. However, 26 other variables are associated with the reservoir and/or each other (green box).

When using variables measured after one year of ART – at the same time as the HIV reservoir was measured (Fig. 3C) – there is little evidence of any correlation with reservoir size or between variables. These data suggest that certain clinical and immunological variables are the key determinants of the HIV DNA level just prior to ART initiation, which is, in turn, the main predictor of HIV reservoir size on ART.

### CD8 T cell activation and memory expansion are the key determinants of HIV DNA levels

To more formally assess the strength and independence of these relationships, regression models were fitted to the data. Correlative analyses reinforced the role for baseline total HIV DNA as the key predictor of reservoir size at one year. For this reason, we concentrated on models to assess which variables were most highly related with baseline total HIV DNA (Fig. 4A).

**Figure 4.**
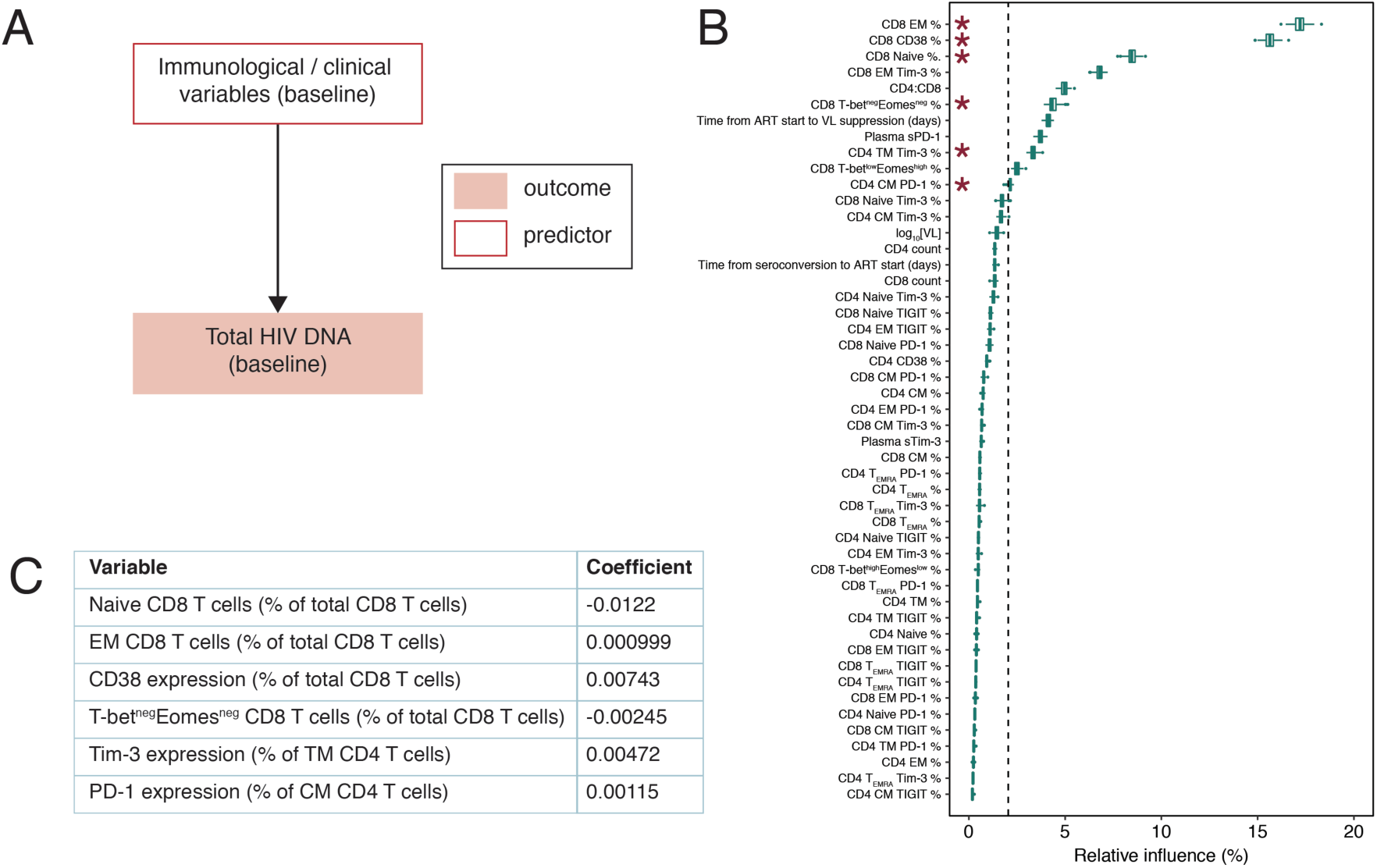
Immunologic and clinical variables which relate to baseline total HIV DNA. **(A)** Structure for models shown in B and C. **(B)** Boosted regression tree model to assess predictors of baseline total HIV DNA (49 predictors, n=62); boxplots show the summary of 100 model runs. Influential predictors were defined as those whose relative contribution was greater than 100 divided by the total number of covariates, this value is indicated by the dashed vertical line. Variables which were also selected in C are marked with an asterisk. **(C)** Least absolute shrinkage and selection operator (LASSO) output for predictors of baseline total HIV DNA (49 predictors, n=59, deviance explained 0.67). Variables which do not significantly contribute to the model have a coefficient of zero; only those with a non-zero coefficient are shown. Missing values were imputed for individuals with some but not all immunologic measures available.

The dataset poses several challenges in the fitting of multivariable regression models, especially as 8.4% of observations are missing (Supplementary Fig. 3) due to unavailable samples, poor sample viability or low cell count. The large number of parameters measured relative to observations, as well as the strong correlations between many of these variables, was also problematic. To ensure the robustness of any conclusions, two different models (boosted regression tree (BRT) and least absolute shrinkage and selection operator (LASSO)) with different approaches to complex data were fitted and their outputs compared.

BRT is a machine learning approach that builds a series of regression trees, with each subsequent tree iteratively aiming to improve the previous fit by focusing on data poorly modelled by the existing set of trees. This approach is able to handle missing data, does not make prior assumptions about the effect of potential predictor variables and can handle high-dimensional interactions (33). A BRT model was fitted with baseline total HIV DNA as the outcome, and all other baseline variables as predictors. The relative importance of each predictor variable included in this BRT model is shown in Fig. 4B. The relative influence of each variable is estimated based on the number of times that variable is selected for splitting and the improvement to the model as a result of that split averaged across all trees; a higher number indicates a greater effect of the variable. We defined influential predictors as those with a relative influence value greater than 100 divided by the total number of variables (indicated by the dashed line). The figure shows that 10 of the 62 predictors had a consistent influence in predicting baseline reservoir size. Notably, CD8 memory subsets (the proportion of EM and naïve cells), as well as CD8 CD38 expression were the variables with the highest relative influence.

LASSO is a multivariable regression designed to cope with multi-collinearity and large numbers of predictors by adding a penalty to the coefficient of each term (with the ability to penalise coefficients to zero) thus performing variable selection. Missing values were imputed using a random forest based method. Linear LASSO models were fitted to the data (Fig. 4C) and selected six variables which were independently predictive of baseline HIV DNA. All six were also selected by the BRT model. The variables with greatest influence on baseline HIV DNA were associated with CD8 memory expansion (the proportion of naïve and EM, as well as T-bet^neg^Eomes^neg^ CD8 T cells) and CD4 memory ICR expression (PD-1 on TM and Tim-3 on CM CD4 T cells), as well as CD38 expression on CD8 T cells.

A sensitivity analysis was conducted by constructing the same model using only observations that were complete (no imputation). The overall results of this model were the same (although due to decreased power fewer variables were selected) and are shown in Supplementary Table 1.

### HIV DNA pre-ART is the dominant predictor of reservoir size at one year on ART

Having established which variables were related to baseline HIV DNA, regression models were then fitted to explore if any of clinical or immunological variables had additional, independent relationships with reservoir size at one year. Total HIV DNA at baseline was the most influential variable when combined with other pre-ART or one year variables (Table 2; Supplementary Fig. 4 shows the corresponding BRT models and Supplementary Table 1 the LASSO models with no imputation).

**Table 2.** Predictors of reservoir size at one year

|  | Model A | Model B |
| --- | --- | --- |
| Predictors included | Baseline clinical and immunological variables (including baseline total HIV DNA) | 1 year immunological variables and baseline total HIV DNA |
|  | *Coefficient* | *Coefficient* |
| Baseline log<sub>10</sub>[total HIV DNA] | 0.339 | 0.311 |
| Time from ART start to VL suppression (days) | 0.000342 | - |

Least absolute shrinkage and selection operator (LASSO) output for predictors of reservoir size (total HIV DNA at 1 year). Model A includes all baseline clinical and immunological variables, including baseline total HIV DNA (50 predictors, n=60, deviance explained 0.58). Model B includes all immunological measures at 1 year along with baseline total HIV DNA (1 year; 44 predictors, n=61, deviance explained 0.47). Variables which do not significantly contribute to the model have a coefficient of zero; only those with a non-zero coefficient are shown. Missing values were imputed for individuals with some but not all immunologic measures available.

No immunological variables measured at one year impacted reservoir size independently of the baseline HIV DNA levels, consistent with the modest correlations observed in the Fig. 3C corrgram (Model B in Table 2, Supplementary Fig. 4B). Of the clinical variables and immunological variables measured at baseline, only ‘time from ART start to VL suppression’ influenced reservoir size independently of the baseline HIV DNA (Model A in Table 2, Supplementary Fig. 4A).

### Reservoir size is related to HLA class I

Although we did not have access to HIV-specific immune responses for HEATHER, participants were typed for HLA class I. These alleles have the strongest consistent relationship with viral control during HIV infection, and HLA type can be considered as a surrogate marker of potential CD8-driven HIV-specific immunity (34, 35). Figure 5 shows the relationship between HLA class I alleles and total HIV DNA at one year post-ART initiation. Alleles associated with viral control (red) can be seen clustering towards the left hand side of the plot associated with low HIV DNA levels. The converse is seen for alleles associated with progression (blue). The same relationship was observed with baseline HIV DNA (data not shown) and supports previous findings in a different PHI cohort (SPARTAC) (7). Together, these data are consistent with HIV specific immunity, the general immune landscape and clinical parameters all contributing to the size of the HIV reservoir on ART.

**Figure 5.**
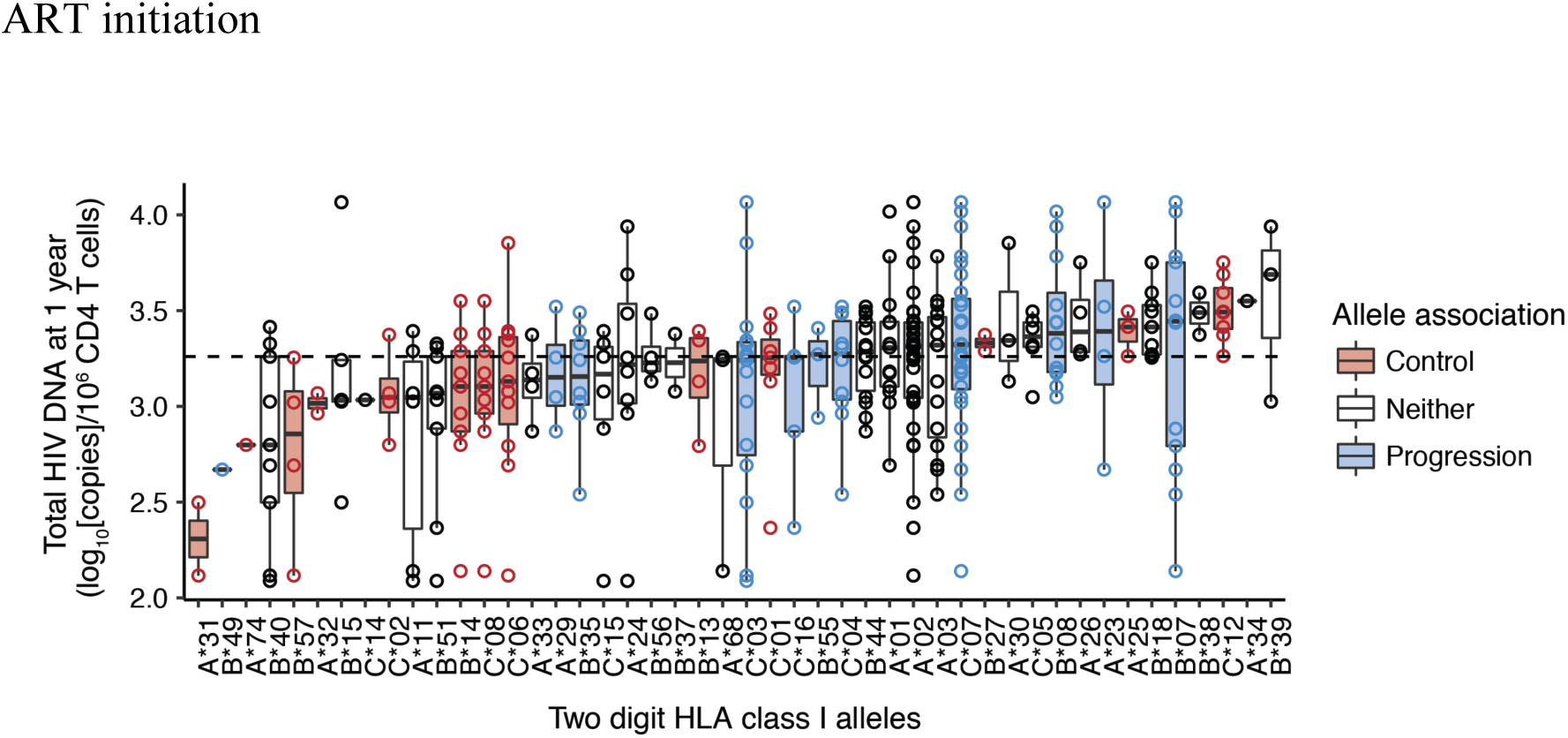
Relationship between HLA class I alleles and total HIV DNA at 1 year post-ART initiation. Relationship between HLA class I alleles and total HIV DNA at 1 year post-ART initiation. Alleles, shown on the x-axis, are ordered based on the median value of total HIV DNA for all individuals possessing that allele. Boxplots show the distribution of total HIV DNA in amongst individuals possessing that allele, and each observation is shown as an open circle. Data is shown for 61 individuals and a total of 333 alleles. For one individual only B alleles were available and are included here; similarly for another individual only A and C alleles were available. Where individuals were homozygous for a given allele, this is only shown once. The dashed line shows the median value of total HIV DNA for the entire cohort. Alleles were classified as being associated with disease progression (blue) or control (red), or neither (white), based on those identified in the International HIV Controllers Study at a significance level of 0.05 (34).

Figure 6 summarises our findings. The only two independent variables that predicted HIV reservoir size after one year of ART were total HIV DNA at pre-therapy baseline and the time taken to achieve viral load suppression after starting therapy. Baseline HIV DNA was associated with HLA class I type and specific markers of T cell activation, expansion and exhaustion.

**Figure 6.**
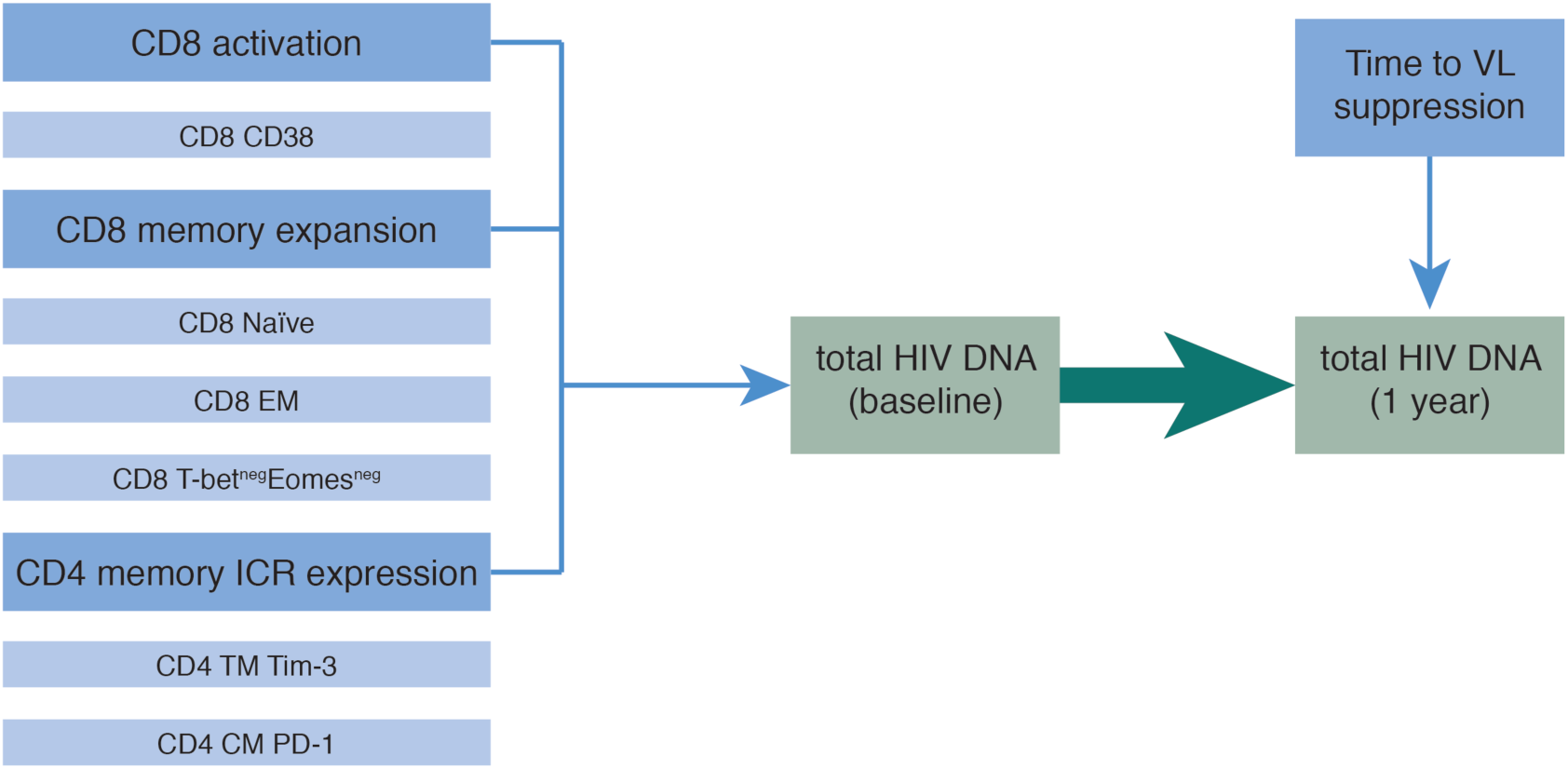
Model for factors which influence reservoir size in treated PHI.

## Discussion

This work demonstrates the importance of immunological events before ART initiation in determining the subsequent reservoir size. The level of HIV DNA prior to ART was the most important predictor of reservoir size a year later, suggesting that reservoir size is “set” early on. Of note, the level of HIV DNA pre-ART was more closely related to CD8 T cell activation, memory expansion and CD4 ICR expression than any clinical parameters, including pre-ART VL. Several of the associations with reservoir size presented here have been observed previously – but generally in isolation. This is the first study to define the independence of relationships between T cell activation, memory expansion and ICR expression with eventual reservoir size.

One of the key findings of this analysis is that the HIV DNA level prior to ART initiation was the most influential predictor of subsequent reservoir size. Several other studies have shown a relationship between pre-therapy HIV DNA levels and those once virologically suppressed on ART (16, 36, 37). These findings also support previous research suggesting that set point HIV DNA levels in untreated individuals are reached shortly after peak VL (38), and extend it to show that this is true even if ART is started early.

Higher levels of initial T cell activation were associated with increased reservoir size. This is consistent with the SPARTAC study where CD38 expression at baseline was significantly associated with total HIV DNA at that time (7). It is possible that this is driven by higher initial viral burden; it could also reflect poorer CD8 effector function suggesting that activation and dysfunction may already be coupled during PHI. A recent study of soluble biomarkers during acute infection demonstrated that several of these were related to HIV reservoir size (also measured by total HIV DNA) prior to and following 96 weeks of ART, independently of VL (17). The soluble biomarkers identified can all be produced by myeloid cells in response to IFN-α/*γ* signalling; the authors speculate that this could indirectly reflect innate and/or T cell responses to viral replication (17). The analysis presented here supports these findings, and directly demonstrates a link between T cell activation and reservoir size.

Cross-sectional studies of primary and chronic HIV infection have shown relationships between contemporaneous HIV DNA and CD8 T cell activation during ART (13, 15, 39), although this has not been consistently demonstrated (37, 40). Such a link between CD8 activation following ART and HIV DNA levels was not observed here. Of note, in this work CD38 expression was used to define activation and most prior studies have used HLA-DR/CD38 co-expression. Work from Cockerham *et al.* suggests that HIV DNA may only relate to HLA-DR and not CD38 expression (15) and could explain this discrepancy.

Gandhi *et al.* observed a relationship between T cell activation during chronic infection and HIV DNA in individuals commencing treatment at this time (16). Unlike in early infection studies ((17) and this analysis), this relationship did not persist following adjustment for baseline VL-suggesting time-dependent differences in the relative influence of baseline VL and T cell activation as predictors.

It is interesting to note that the VL was only modestly correlated with baseline HIV DNA, as we had hypothesised that these would be closely linked. Indeed, VL was not a significant predictor here. Several studies have demonstrated a relationship between pre-therapy VL and HIV DNA (16, 37, 41-44). Notably, many of these studies included individuals treated during chronic infection, where the VL will have reached set point. In contrast, during PHI the viral load is substantially more labile. Within this cohort there was a link between baseline VL and the time after estimated seroconversion that this was measured (Fig. 1E), whereby individuals with more recent seroconversion had a higher VL. This has also been observed in another PHI cohort with similar time since infection (45) and suggests that VL measures taken over this time capture the decline from peak, rather than a steady state. Within HEATHER there is some variation in the time interval between when baseline VL and HIV DNA were measured (median 16 days; range 0-82 days). Both of these factors may contribute to the relatively weak relationship between VL and baseline HIV DNA observed here.

It is also possible that the overall pre-ART viral burden (both duration and magnitude of viraemia) influences HIV reservoir size, and is poorly captured by a single measurement during PHI. Indeed, the consistent observation that earlier ART limits reservoir size, even over the short time periods between the first three Fiebig stages (8, 36) implies a role for the total pre-therapy viral burden. Two findings here suggest an influence of viral burden on overall reservoir size. The first is the previously reported relationship between HLA class I alleles and HIV reservoir size (7), which was confirmed. The well-characterised relationship between HLA class I alleles and VL is mediated via CD8 T cell killing of virally infected cells; stronger CD8 responses could result in smaller HIV reservoir size by both limiting seeding (via VL reduction) and contributing to decay of the pool of infected cells. The second relevant finding is the observation that time to viral load suppression had an influence on reservoir size independently of baseline HIV DNA level. This may be because individuals with longer time to VL suppression may have a window following ART initiation for reservoir seeding to continue. Alternatively, slower time to viral suppression may reflect higher pre-therapy viral burden not captured in the baseline VL measurement. Whilst most individuals commenced ART at or shortly after their baseline visit (median 0 days, 82% within 1 week), a small proportion had a larger interval between these (maximum 48 days) providing additional time for reservoir seeding not captured in the baseline HIV DNA measurement.

It is notable that the two measures of ICR expression which related to reservoir size were measured on TM and CM CD4 T cells, the subsets most enriched for proviral DNA (19-22). Several cross-sectional studies have shown a relationship between PD-1 expression on bulk CD4 T cells during ART and overall reservoir size (13, 14, 18). CM and TM CD4 T cells expressing PD-1 are also enriched for proviral DNA (18, 19). The findings presented here demonstrate a relationship between PD-1 expression on CM CD4 T cells supporting these prior observations and showing that this relationship exists during untreated PHI. Contrasting with prior work, here this relationship did not continue during ART. It is possible that previous studies, which have measured PD-1 on bulk T cells (13, 14, 18), are actually capturing this shift in memory proportions. This discrepancy could also be due to differences between individuals who commence ART in primary, compared with chronic, infection.

Whilst a comprehensive analysis of ICR expression, activation and T cell differentiation was performed, one limitation of this work is that reservoir size was only measured using total HIV DNA. This measure is clinically relevant as lower levels have been associated with delayed viral rebound following treatment interruption (4). Much of the HIV reservoir, however, is not replication competent (46, 47). Whilst total HIV DNA has been suggested to reflect replication-competent reservoir size in cohort studies (48), this work has not assessed if there is any impact of these immunological measures on the quality of proviruses comprising the reservoir. A major strength of this study is its longitudinal design. It is possible, however, that 1 year of follow up is not sufficient time for ART to stabilise the HIV reservoir and that these results may not reflect the eventual long-term reservoir. For individuals for whom samples were available, we have shown that HIV DNA levels one year following ART initiation correlated with those at three years, suggesting the validity of this approach. Several other studies have shown relationships between HIV DNA levels at early time points and those much later after ART initiation (16, 37, 49), further supporting the use of this time point here.

This work has shown that the magnitude of the early immunological insult, reflected in CD8 T cell activation and memory expansion drives HIV reservoir size. These results suggest that targeting of host or viral factors which lead to early viral expansion and T cell activation may be a way of limiting HIV reservoir size, and confirm the importance of starting ART as early as possible.

## Methods

### Participant information

HEATHER is a prospective observational cohort study of individuals who commence ART (and remain on uninterrupted therapy) within 3 months of the date of HIV diagnosis during PHI. Individuals are considered to have PHI if they meet any of the following criteria: HIV-1 positive antibody test within 6 months of a HIV-1 negative antibody test, HIV-1 antibody negative with positive PCR (or positive P24 Ag or viral load detectable), RITA (recent incident assay test algorithm) assay result consistent with recent infection, equivocal HIV-1 antibody test supported by a repeat test within 2 weeks showing a rising optical density or having clinical manifestations of symptomatic HIV seroconversion illness supported by antigen positivity. The time of seroconversion was estimated as the midpoint between the most recent negative or equivocal test and the first positive test for those who met relevant criteria, the date of RITA test minus 120 days and as the date of test for all other participants. Individuals with co-existent hepatitis B or C infection are not eligible for inclusion in HEATHER. Individuals for inclusion in this analysis were selected at random and based on sample availability.

Cryopreserved peripheral blood mononuclear cells (PBMCs) were used from the closest sample to seroconversion (baseline) and from a sample 9–15 months after commencement of ART (1 year). CD4 count, CD8 count and VL were measured as part of routine clinical care with baseline CD4 and CD8 counts defined as the earliest value prior to the initiation of ART. Similarly, the baseline VL was taken as the earliest value prior to the initiation of ART. In both cases, usually only one value was available.

### Flow cytometry

Cryopreserved PBMCs were thawed in RPMI-1640 medium supplemented with 10% FBS, L-glutamine, penicillin and streptomycin (R10) containing 2.7 Kunitz units/ml of DNAse (Qiagen). Cells were stained in BD Horizon Brilliant Stain Buffer (BD) containing all antibodies and Live/Dead Near IR at 1 in 300 dilution (Life Technologies) at 4°C for 30 minutes.

Panel 1-PBMCs were stained with the following antibodies: CD3 Brilliant Violet (BV) 570 (UCHT1), CCR7 Pacific Blue (G043H7), CD27 AlexaFluor700 (M-T271)[all BioLegend], CD4 BV605 (RPA-T4), CD8 BV650 (RPA-T8)[all BD], PD-1 PE-eFluor610 (eBioJ105), CD45RA FITC (HI100), TIGIT PerCP-eFluor710 (MBSA43)[all eBioscience] and Tim-3 PE (344823)[R&D].

Panel 2-PBMCs were stained with the following antibodies: CD3 BV570, CD38 AlexaFluor700 (HB-7) [BioLegend], CD4 BV605, CD8 BV650, PD-1 PE-eFluor610 and Tim-3 PE. Following this, cells were washed twice prior to fixation and permeabilisation with Foxp3 Buffer Set (BD) as per manufacturer’s directions with reduced volumes to facilitate staining in 96-well plates. Staining for intracellular epitopes was performed at room temperature for 30 minutes in PBS containing 0.5% bovine serum albumin (BSA) and 0.1% sodium azide with the following antibodies: T-bet FITC (4B10)[BioLegend] and Eomes eFluor660 (WD1928)[eBioscience].

For both panels, cells were washed twice, fixed with 2% formaldehyde for 30 minutes, then washed twice and resuspended in phosphate buffered saline (PBS) for acquisition.

All samples were acquired on a LSR II (BD) the day after staining. The same machine was used for all experiments with daily calibration with Cytometer Setup & Tracking beads (BD) to maximise comparability between days. Rainbow Calibration Particles (BioLegend) were also used for cohort phenotyping to minimise batch-to-batch variability. Data were analysed using FlowJo Version 10.8.0r1 (Treestar).

### Soluble PD-1 and Tim-3 quantification

The concentration of sPD-1 and sTim-3 was measured in thawed plasma samples by enzyme linked immunosorbent assay (ELISA) using Human PD-1 (PDCD1) ELISA kit [EHPDCD1] (Thermo Fisher Scientific, Waltham, MA USA) and Quantikine ELISA Human TIM-3 Immunoassay kit [DTIM30] (R&D Systems, Minneapolis, MN USA) as per manufacturers’ instructions at 1:2 and 1:5 dilutions respectively.

### Total HIV DNA quantification

HIV DNA was quantified relative to cell number using qPCR as previously described (50). In brief, cryopreserved PBMCs were thawed (as above) and CD4 T cells were isolated by negative selection using the EasySep Human CD4 Enrichment Kit (Stemcell Technologies) before DNA extraction with the QiaAMP Blood Mini Kit (Qiagen). Cell copy number was initially quantified using an albumin qPCR. 25,000 cell equivalents (and no less than 10,000 cell equivalents) of DNA was then used in a total HIV DNA qPCR, performed in triplicate. The mean number of copies of DNA was normalised to cell number and expressed as copies/10^6^ CD4 T cells.

### HLA-typing

HLA typing was performed to intermediate resolution using PCR with sequence specific primers (PCR-SSP).

### Statistics

Analyses were performed using R (v3.2.2 or v3.4.3) and GraphPad Prism (v7.0b). Except where otherwise specified, p-values <0.05 were considered statistically significant. Simple comparisons were performed using parametric or non-parametric tests as appropriate and are described alongside the results.

Corrgrams were generated using the package corrplot (v0.84). Where some data were missing, pairwise complete observations were used to calculate the correlation coefficients.

Boosted regression tree models were fitted with a Gaussian outcome using the package gbm3 (v2.2). Models were fitted with an interaction depth of 5 with a minimum of 5 observations in a terminal node. The shrinkage parameter was adjusted between 0.0001-0.001 to aim for optimal number of trees to fall in the range of 3000-10000. The optimal number of trees was determined using 10 fold cross-validation, with the number of trees that minimised cross-validation error chosen. Results presented are summarised outcomes of 100 models.

LASSO models (51) were fitted using the R package glmnet (v2.0-16) (52). Gaussian regression models were fitted with an additive linear model (no interactions). The tuning parameter λ was determined using 10-fold cross-validation, with the λ value used being that which minimised cross-validation error plus one standard error. Where data were imputed this was performed using the package MissForest (53), which employs a random forest based method for multiple imputation and has been shown to be superior to other multiple imputation methods in biological datasets with comparable levels of missingness (53, 54).

### Study approval

Recruitment for the HEATHER cohort was approved by the West Midlands-South Birmingham Research Ethics Committee (reference 14/WM/1104). All participants have given informed consent for their participation in these studies.

## Author contributions

GEM, JFr devised the study, analysed data and wrote the manuscript; FS and JH advised on the analysis of data; GEM, MP, EZ, JT, HB, NR, EH and NO performed laboratory analyses; JM managed the patient cohorts and collated demographic data; JT, JL, NN, JFo and SF managed clinical sites and recruitment; NN, JFo, SF, CBW, JFr led on study design and management.

## Acknowledgments

We thank the participants of HEATHER. The HEATHER study is conducted as part of the CHERUB (Collaborative HIV Eradication of Reservoirs: UK BRC) collaboration. CHERUB Steering Committee: Andrew Lever (University of Cambridge), Mark Wills (University of Cambridge), Jonathan Weber (Imperial College, London), Sarah Fidler (Imperial College, London), John Frater (University of Oxford), Lucy Dorrell (University of Oxford), Mike Malim (King’s College, London), Julie Fox (King’s College London), Ravi Gupta (University College London), Clare Jolly (University College London). Thank you to the following who have been involved with the recruitment of HEATHER at the trial sites. Kristin Kuldanek, Heather Lewis, Rebecca Hall (St Mary’s Hospital), Teresa Solano (St Thomas’ Hospital), Sathya Visvendra, Rhian Bull and Gabriele Pasluostaite (Chelsea & Westminster Hospital).

## Supplementary Materials

Supplementary Figure 1 – Viral load sampling frequency

Supplementary Figure 2 – Reservoir size at 3 years post-ART initiation

Supplementary Figure 3 – Missing immunological and clinical data

Supplementary Figure 4 – Boosted regression tree results to assess the relative influence of predictors of reservoir size at 1 year

Supplementary Table 1 – LASSO models using data without imputation

