## Supplementary data for "HIV reservoir size is determined prior to ART initiation and linked to CD8 T cell activation and memory expansion"

**Supplementary Figure 1.** Viral load sampling frequency

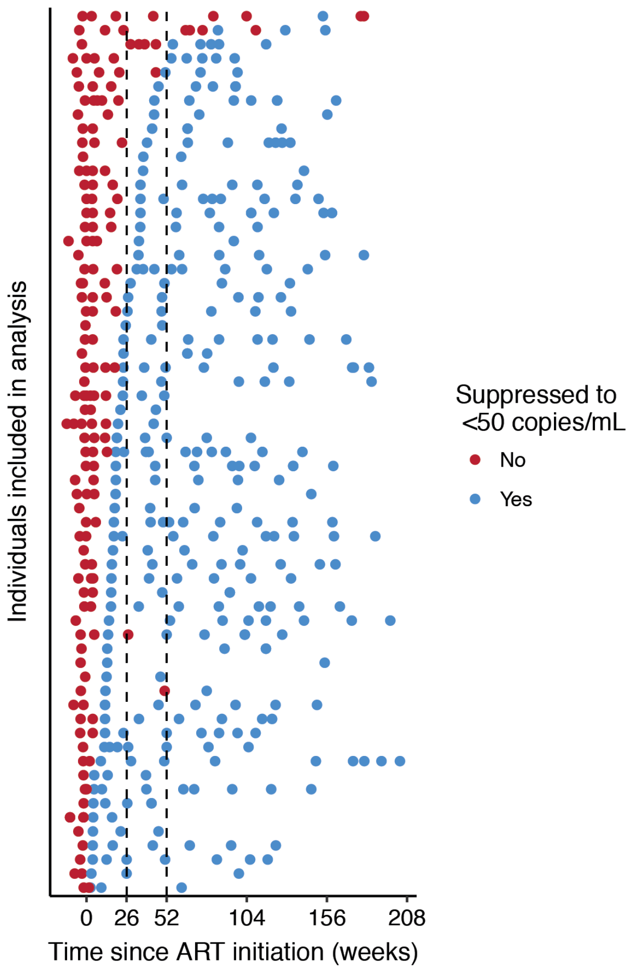


The viral load sampling frequency in individuals (n=63) included in the analysis. Each VL measurement is plotted at the time following ART initiation that it was taken and coloured according to whether the VL value was suppressed to <50 copies/mL (red if not suppressed, blue if suppressed). Individuals are ordered based on the time to viral load suppression (first measurement <50 copies/mL).

**Supplementary Figure 2.** Reservoir size at 3 years post-ART initiation

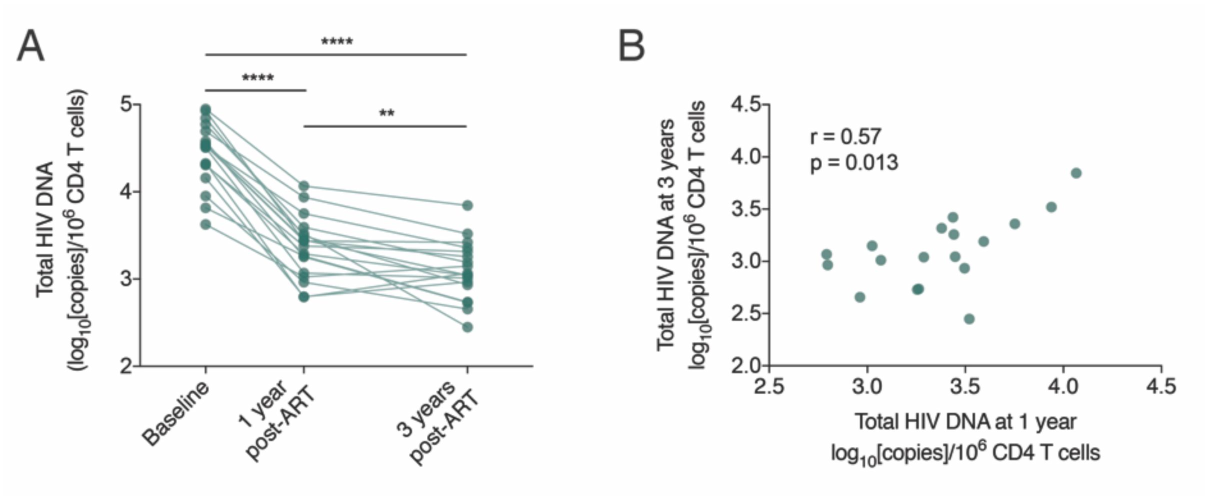


Relationship between total HIV DNA measured at baseline, 1 year following ART initiation and 3 years following ART initiation for a subset of individuals (n=18). For **(A)** comparison was made using a ANOVA (with pairing; overall $p<0.0001$) with post-hoc testing using Holm-Sidak's multiple comparison test. ** indicates $p <0.01$, **** indicates $p<0.0001$. **(B)** a Pearson's correlation was performed.

**Supplementary Figure 3.** Missing immunological and clinical data

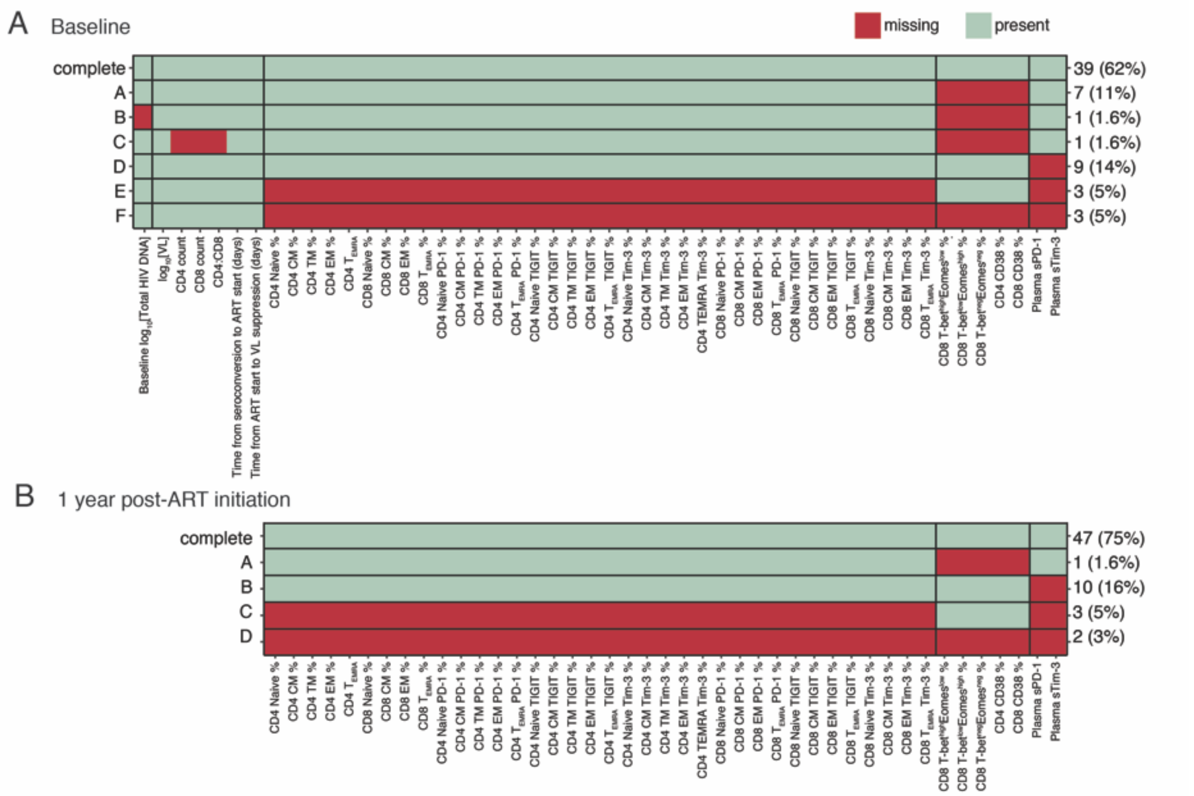


Missing immunological and clinical data. This diagram shows the patterns of missing data at baseline **(A)** and following 1 year of ART **(B)** for the 63 individuals included in analyses presented. Each row represents a pattern of data missingness and the right hand side shows how many individuals have this pattern. For some individuals there are no immunological observations available at a given time point. These are Pattern F (n=3) at baseline (all three samples had very low viability upon thawing and were not able to be included) and Pattern D (n=2) at 1 year (these individuals had not returned for 1 year visits at the time the assays were performed).

**Supplementary Figure 4.** Boosted regression tree results to assess the relative influence of predictors of reservoir size at 1 year

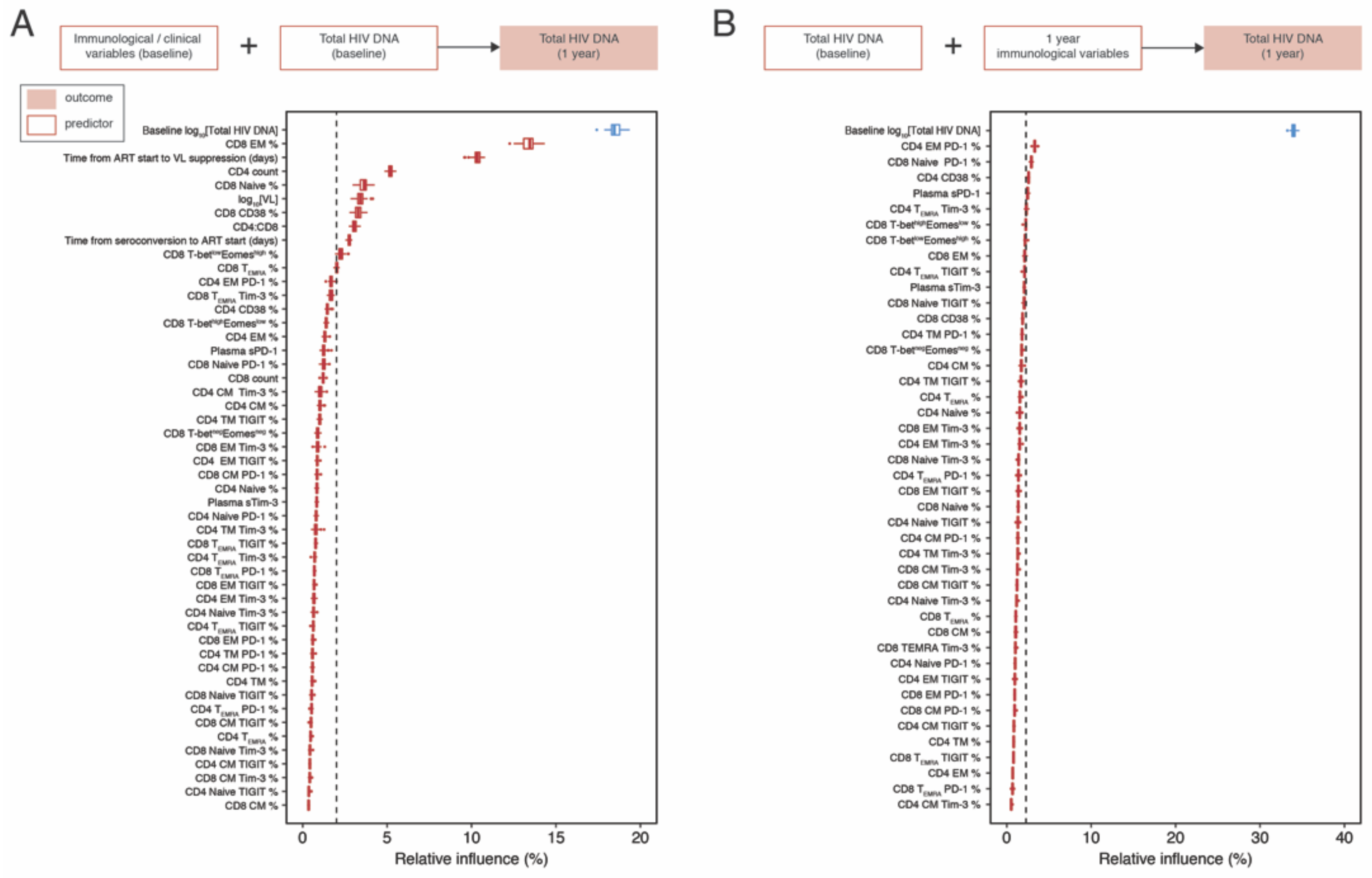


Boosted regression tree models to assess the relative influence of predictors of reservoir size (1 year total HIV DNA). **(A)** Includes all baseline clinical and immunological variables (50 predictors, n=63). **(B)** Includes all immunological measures at 1 year (1 year; 44 predictors, n=63). In both models, total HIV DNA at baseline was included as a predictor and is highlighted in blue. Boxplots show the summary of 100 model runs. Influential predictors were defined as those whose relative contribution was greater than 100 divided by the total number of covariates, this value is indicated by the dashed vertical line.

**Supplementary Table 1.** LASSO models using data without imputation

| Corresponding model | Fig. 4C | Table 2 – Model A | Table 2 – Model B |
|---|---|---|---|
| Outcome | Baseline total HIV DNA | 1 year total HIV DNA | 1 year total HIV DNA |
| Predictors included | Baseline clinical and immunological variables | Baseline clinical and immunological variables | 1 year immunological variables |
| n | 39 | 39 | 46 |
| Deviance explained | 0.64 | 0.51 | 0.42 |
| | | *Coefficient* | *Coefficient* |
| Naive CD8 T cells (% of total CD8 T cells) | -0.0157 | - | - |
| CD38 expression (% of total CD8 T cells) | 0.00620 | - | - |
| Tim-3 expression (% of TM CD4 T cells) | 0.00242 | - | - |
| Baseline $log_{10}$[total HIV DNA] | N/A | 0.271 | 0.248 |
| Time from ART start to VL suppression (days) | - | 0.000458 | N/A |

Least absolute shrinkage and selection operator (LASSO) output for predictors of total HIV DNA. LASSO models presented here were constructed excluding any observations with missing variables (complete cases only) rather than imputing these missing values. Variables which do not significantly contribute to the model have a coefficient of zero; only those with a non-zero coefficient are shown. N/A means that variable was not included in model construction.
